# Yellow Fever Emergence: Role of Heterologous Flavivirus Immunity in Preventing Urban Transmission

**DOI:** 10.1101/2024.03.03.583168

**Authors:** Divya P. Shinde, Jessica A. Plante, Dionna Scharton, Brooke Mitchell, Jordyn Walker, Sasha R. Azar, Rafael K. Campos, Lívia Sacchetto, Betânia P. Drumond, Nikos Vasilakis, Kenneth S. Plante, Scott C. Weaver

## Abstract

During major, recent yellow fever (YF) epidemics in Brazil, human cases were attributed only to spillover infections from sylvatic transmission with no evidence of human amplification. Furthermore, the historic absence of YF in Asia, despite abundant peridomestic *Aedes aegypti* and naive human populations, represents a longstanding enigma. We tested the hypothesis that immunity from dengue (DENV) and Zika (ZIKV) flaviviruses limits YF virus (YFV) viremia and transmission by *Ae. aegypti*. Prior DENV and ZIKV immunity consistently suppressed YFV viremia in experimentally infected macaques, leading to reductions in *Ae. aegypti* infection when mosquitoes were fed on infected animals. These results indicate that, in DENV- and ZIKV-endemic regions such as South America and Asia, flavivirus immunity suppresses YFV human amplification potential, reducing the risk of urban outbreaks.

**One-Sentence Summary:** Immunity from dengue and Zika viruses suppresses yellow fever viremia, preventing infection of mosquitoes and reducing the risk of epidemics.

## Introduction

The conspicuous absence of yellow fever (YF) in highly vulnerable regions of Asia has been an enigma for over a century *(1)*. YF, a mosquito-borne disease that can be hemorrhagic with a case-fatality rate of 20-50%, is caused by YF virus (YFV) (*Flaviviridae, Orthoflavivirus*) (2). Although a vaccine is available *(3)* manufacturing limitations and shortages have occurred in the face of outbreaks, raising concerns regarding the ability to control future epidemics (*4)*.

YFV circulates in two ecologically and evolutionarily distinct transmission cycles *(5)*. In the urban or epidemic cycle, transmission involves human amplification and the peridomestic mosquito vector, *Aedes (Stegomyia) aegypti*. This cycle was responsible for a recent epidemic in Angola *(6)*, which subsequently spread to the Democratic Republic of Congo *(7)*. In the sylvatic or enzootic cycle, YFV is transmitted among non-human primates (NHPs) by forest mosquitoes in South America and Africa *(8,9*). In the rural zone of emergence bordering forests, the virus can spill over from enzootic vectors to both NHPs and humans. Increased human encroachment in these regions with high densities of enzootic vectors can result in epidemics, as occurred in Brazil from 2016- 2019 *(8)*. However, despite being the largest sylvatic outbreak reported since 1942 *(10*) and an abundance of *Ae. aegypti*, there were no reports of human-amplified urban transmission cycle *(8*). Although vaccination coverage varies across Brazil *(11)*, YFV infections during this outbreak included areas outside the recommended regions, where low herd immunity provided an ideal opportunity for *Ae. aegypti*-borne transmission. Thus, the lack of an urban cycle in Brazil since 1942 and the complete historic lack of YF in Asia, despite completely naïve populations and abundant vectors, remains a mystery *(1*).

Several theories have been proposed to explain the lack of YF in Asia *(1)*: 1) A paucity of introductions from Africa. This seems unlikely because chikungunya (CHIKV) and Zika (ZIKV) viruses, which also originated in sub-Saharan Africa and are transmitted by *Aedes* mosquitoes, have long histories of spread to establish urban transmission in Asia *(12*). Furthermore, the well-documented introduction of YFV via travelers from Angola to China in 2016 failed to establish autochthonous transmission *(1,13*); 2) Asian and South American *Ae. aegypti* strains are less competent vectors compared to African strains. However, laboratory experiments showed that *Ae. aegypti* populations from various Asian and South American countries are capable of YFV transmission *(14–*16); 3) Another theory suggests that pre-existing immunity from related flaviviruses, particularly dengue virus (DENV) with very high seroprevalence in Asia, could act as an immune barrier to prevent the establishment of human-amplified YFV transmission (*17*,18). Supporting this hypothesis, a previous study showed that dengue-immune monkeys exhibit lower viremia in response to experimental South American and African YFV infection, compared to dengue-naïve animals *(17*). Other studies in rhesus macaques and vervet monkeys suggest a partial protective role of previous flavivirus immunity against YFV infection, particularly in reducing viremia (19). However, the effect of flavivirus immunity on prevention of *Ae. aegypti* infection has not been addressed.

In addition to longstanding DENV circulation in South America and Asia, ZIKV caused a major epidemic in the Americas from 2014 to 2016, infecting over 50% of many neotropical human populations *(20)*. The peak of this outbreak preceded the 2016 YF epidemic, raising the question of whether preexisting ZIKV immunity may also have influenced YFV infection dynamics including viremia, which could influence interhuman mosquito transmission and thus human exposure. Since ZIKV has been endemic to Asia for many decades (*2*1), its immunity could also have reduced the risk of YF there.

We tested the hypothesis that heterologous flavivirus immunity from DENV and ZIKV infection modulates YFV infection of humans, reducing viremia and mosquito infection, thereby rendering people less competent as amplification hosts to sustain urban transmission. We used an NHP model with heterologous flavivirus immunity by assessing viremia, markers of YF disease, and infection of *Ae. aegypti* during periods of peak viremia. Our findings revealed that prior immunity to DENV and ZIKV generally reduces YFV viremia, as well as mosquito infection and subsequent transmission potential. Heterologous immunity also reduces the elevation of liver transaminases in most animals, suggesting protection against hepatic disease.

## Results

### Dengue virus (DENV-2) and ZIKV-immune non-human primates (NHPs) have modulated YFV viremia

We examined the effect of pre-existing DENV-2 and ZIKV immunity on YFV infection of cynomolgus macaques (Figure 1a). The initial infections were described in a previous study *(22)*; 23 animals were exposed to *Ae. aegypti* mosquitoes sham-infected with PBS or infected with one of four virus strains: DENV-2 P8-1407 or ZIKV DakAr 41525 (sylvatic strains), DENV-2 NGC or ZIKV PRVABC59 (endemic/epidemic strains). In our study, 6-9 months after primary exposure, all animals were challenged with a Brazilian YFV strain (YFV_BR_MG-2017NHP15, hereafter YFV BR17) isolated in 2017, during the major YF epidemic. Prior to YFV challenge, sera were assayed by foci-reduction neutralization test (FRNT) for antibodies against all three viruses. Flavivirus-naïve (exposed to uninfected mosquitoes) animals were seronegative for all four serotypes of DENV, as well as ZIKV, and YFV (Figure 1b, 1c, Supplementary Table 1). Of the 10 DENV-2 P8-1407-exposed animals, eight had neutralizing antibodies against DENV-2, whereas two animals were seronegative by FRNT (<20; Figure 1b). All animals exposed to DENV- 2 NGC and ZIKV DakAr 41525 had homologous neutralizing antibodies (Figure 1b, 1c). One of the four animals exposed to ZIKV PRVABC59-infected mosquitoes did not seroconvert by FRNT (Figure 1c).

**Fig 1:**
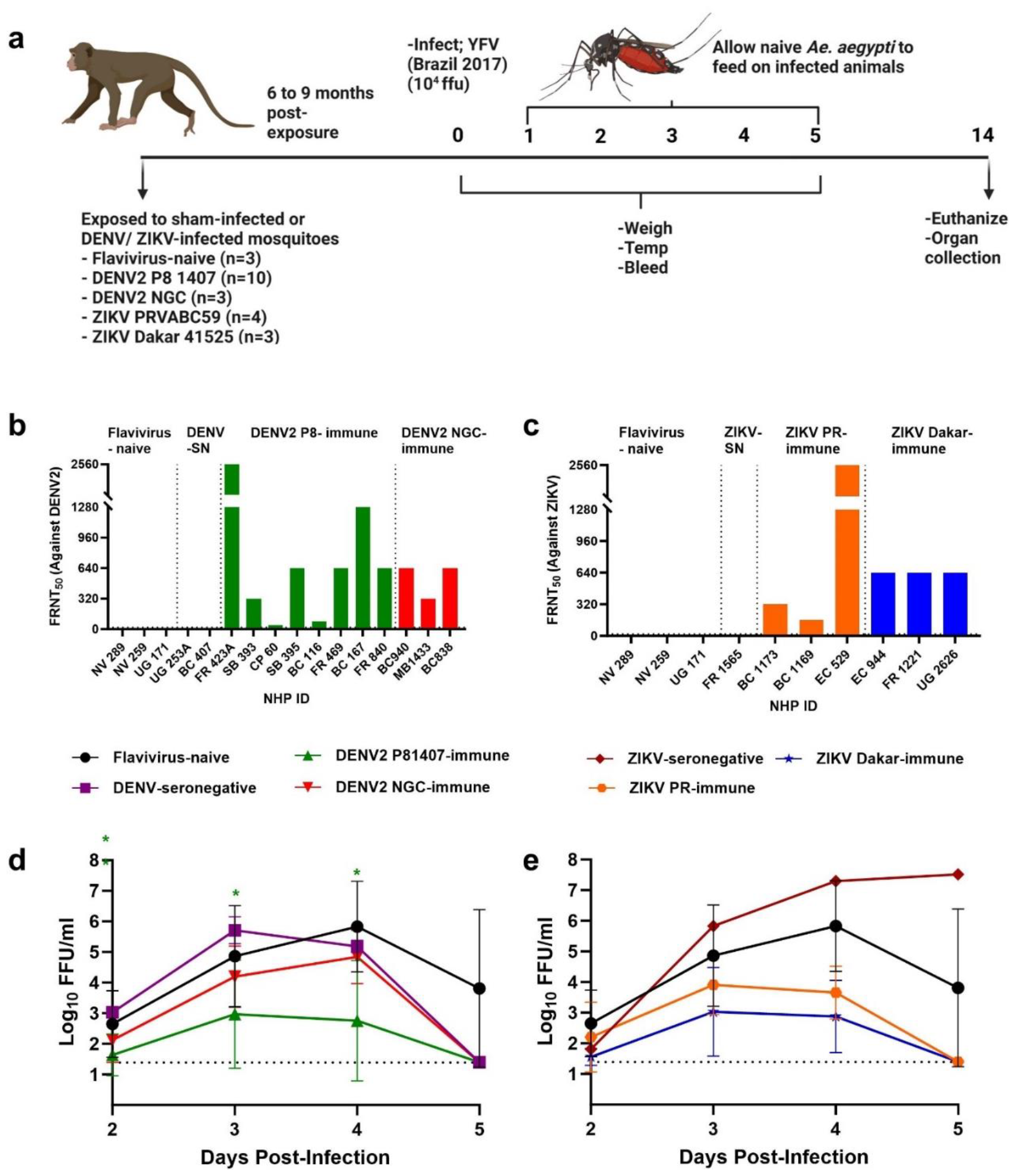
DENV-2 and ZIKV-immune NHPs have modulated YFV viremia. **(A)** Experimental design in cynomolgus macaques (created with BioRender.com). **(B)** Neutralizing antibody titer (FRNT_50_) against DENV-2 in flavivirus-naïve and DENV-2-exposed animals, prior to YFV challenge. **(C)** Neutralizing antibody titer against ZIKV in flavivirus-naïve and ZIKV-exposed animals. Animals with homologous FRNT_50_ titers below the limit of detection (1:20) were reclassified as seronegative (SN). YFV viremia levels in flavivirus-naïve and DENV-2-exposed **(D)** and ZIKV-exposed **(E)** animals from 2-5 DPI. Viremia levels were measured by focus-forming assays. Log_10_ transformed viremia values were analyzed by two-way ANOVA with Tukey’s multiple comparison test. A significant difference in DENV-2 P8-immune and DENV-SN groups was observed at 2 (p=0.0039), 3 (p=0.0203), and 4 (p=0.0398) DPI. For all graphs, symbols represent the mean, and error bars represent the standard deviation. Statistical significance (*) color is attributed to the significant difference between that group compared to the flavivirus-naïve group. Reported p-values are based on the results of the respective post-hoc tests. *p<0.05, ** p<0.01.

Upon subcutaneous YFV infection with a dose of 10^4^ focus-forming units (FFU), DENV-2- and ZIKV-FRNT-positive animals had lower levels and shorter duration of viremia compared to flavivirus-naïve animals (Figure 1d and 1e, Supplementary Table 2, 3). The two DENV- seronegative animals (fed upon by infected mosquitoes but without FRNT seroconversion) had comparable viremia levels to flavivirus-naïve animals. However, the ZIKV-seronegative animal (NHP ID: FR1565, exposed to the bite of an infected mosquito but FRNT-negative) had higher titered and longer viremia compared to the flavivirus-naïve animals (Figure 1e). Surprisingly, unlike any other YFV-infected animal, FR1565 was laterally recumbent and had to be euthanized by day 9 post-challenge. No visible signs of illness were observed in any other animals.

To investigate further if FRNT-seronegative animals bitten by infected mosquitoes became infected with DENV or ZIKV, we performed ELISA to detect binding but non-neutralizing antibodies. Interestingly, both of the DENV-2 P8-1407-seronegative and the one ZIKV- seronegative animal had low levels of IgG antibodies, suggesting that infection did occur despite the lack of neutralizing antibody induction. Due to extensive cross-reactions amongst flaviviruses, non-neutralizing antibodies were not measured against heterologous viruses.

### Mosquitoes fed on YFV-viremic NHPs with previous DENV and ZIKV immunity are refractory to YFV infection

The gold standard measure of human amplification competence is the ability of viremic people to infect *Ae. aegypti*, leading to efficient epidemic transmission. To mimic natural infection, we allowed naïve female *Ae. aegypti* mosquitoes, derived from a colony originating in Salvador, Brazil, to feed on infected animals 1-5 days post-YFV infection; 100 mosquitoes were exposed to each animal per day. Fully engorged mosquitoes were then incubated under tropical conditions (28°C, 75% relative humidity) for 14 days. Mosquito bodies, legs, and saliva were collected to represent initial infection, disseminated infection, and transmission potential, respectively. Most mosquitoes (bodies) that fed on flavivirus-naïve animals had high rates of YFV infection on peak viremia days 3 (Figure 2a) and 4 (Figure 2b). In contrast, most mosquitoes that fed on DENV- and ZIKV-immune animals were refractory to infection (Supplementary Table 4). Infection rates of mosquitoes that fed on flavivirus-naïve animals were significantly higher than those that fed on DENV-2 NGC-, DENV-2 P8-1407-, ZIKV PRVABC-, and ZIKV DakAr 41525-immune animals (p<0.0001; Supplementary Table 5). Significant differences were not observed between mosquitoes (bodies) that fed on flavivirus-naïve animals versus DENV-2-seronegative animals at 3 days post-infection (DPI) (p=0.2678), and the ZIKV seronegative animal at 4 DPI (p=0.3132). Viral titers in mosquito bodies were significantly higher in flavivirus-naïve versus -immune NHPs (Figure 2c, 2d; p<0.0001). Dissemination from the digestive tract (YFV-positive legs) was observed primarily in the flavivirus-naïve and -seronegative groups (Supplementary Table 4) and transmission potential (YFV-positive saliva) was observed only for a mosquito that fed on a flavivirus-naïve animal.

**Fig 2:**
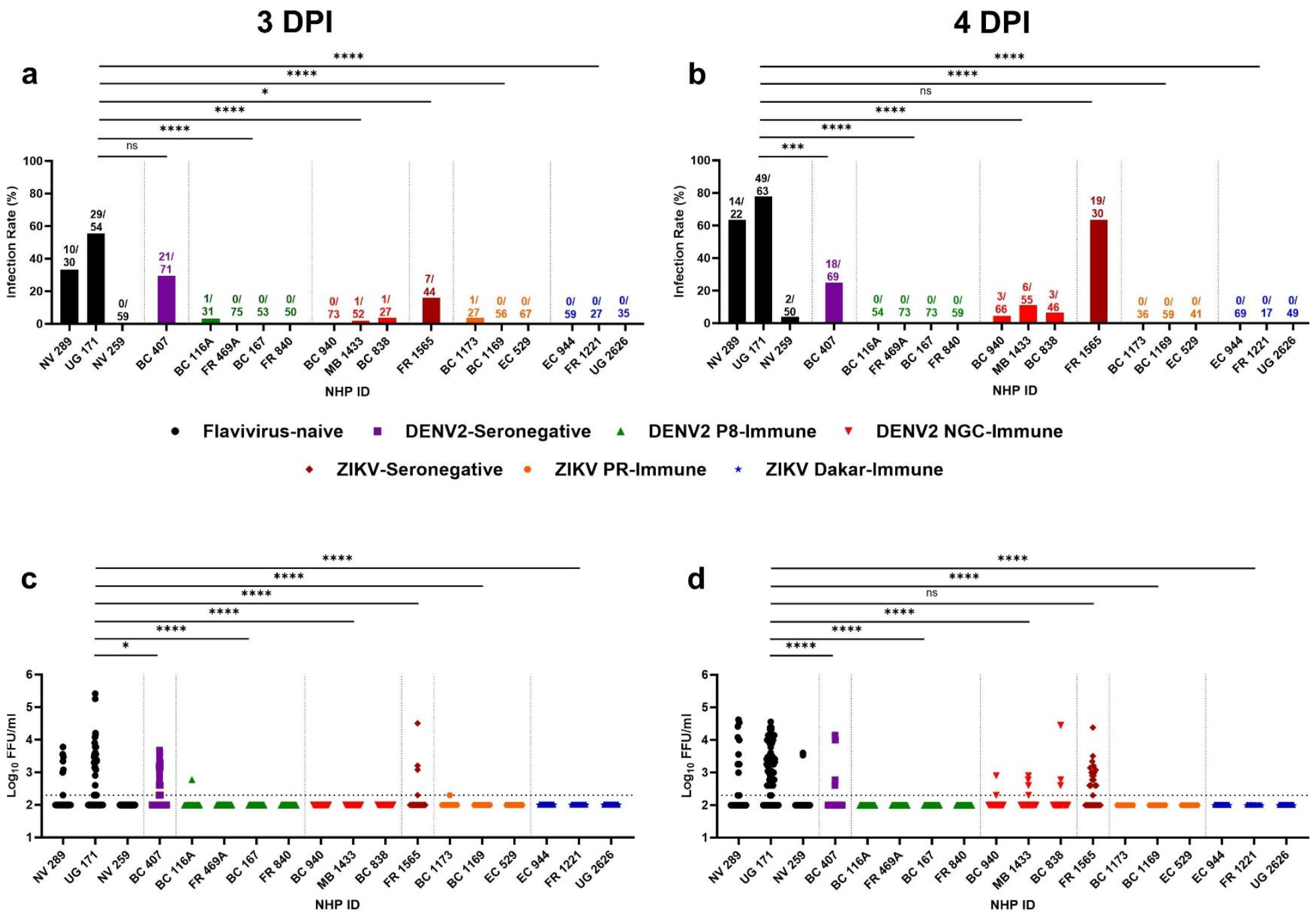
Majority of *Ae. aegypti* fed on YFV-infected DENV-2 and ZIKV-immune NHPs are refractory to YFV infection. Mosquitoes fed on infected NHPs were allowed to incubate for 14 days at 28°C and 75% humidity. Mosquito infection rate (number of mosquito bodies infected/total number of engorged mosquitoes) is shown for mosquitoes fed on NHPs at day 3 **(A)** and day 4 **(B)** post-YFV infection. Each bar represents a cohort of mosquitoes fed on individual NHPs at respective time points. Values on top of each bar represent the number of mosquitoes tested positive/ total number of mosquitoes engorged. Log_10_ transformed viral titers in individual mosquito bodies at day 3 **(C)** and day 4 **(D)** post-YFV infection. Mosquito infectivity and viral titers have been shown for each animal since there is a variation in viremia levels despite a similar immune status. For statistical analyses, the data for each group was combined and compared with the outcomes for the flavivirus-naïve group by two-tailed Fisher’s exact test. Similarly, viral titers have been shown for group of mosquitoes that fed on individual NHPs. For statistical analyses, data from each group was combined and Log_10_ transformed values were analyzed by one-way ANOVA with Dunnett’s multiple comparison test. *p<0.05, ** P<0.01, ***p<0.001, ****p<0.0001.

Comparing the mosquito infectivity for flavivirus-naïve and DENV-2-immune groups, we observed that infectivity was 20-30% lower for DENV-immune compared to flavivirus-naïve animals with similar YFV viremia levels (5.3 Log_10_ FFU/ml) (Figure 3).

**Fig 3:**
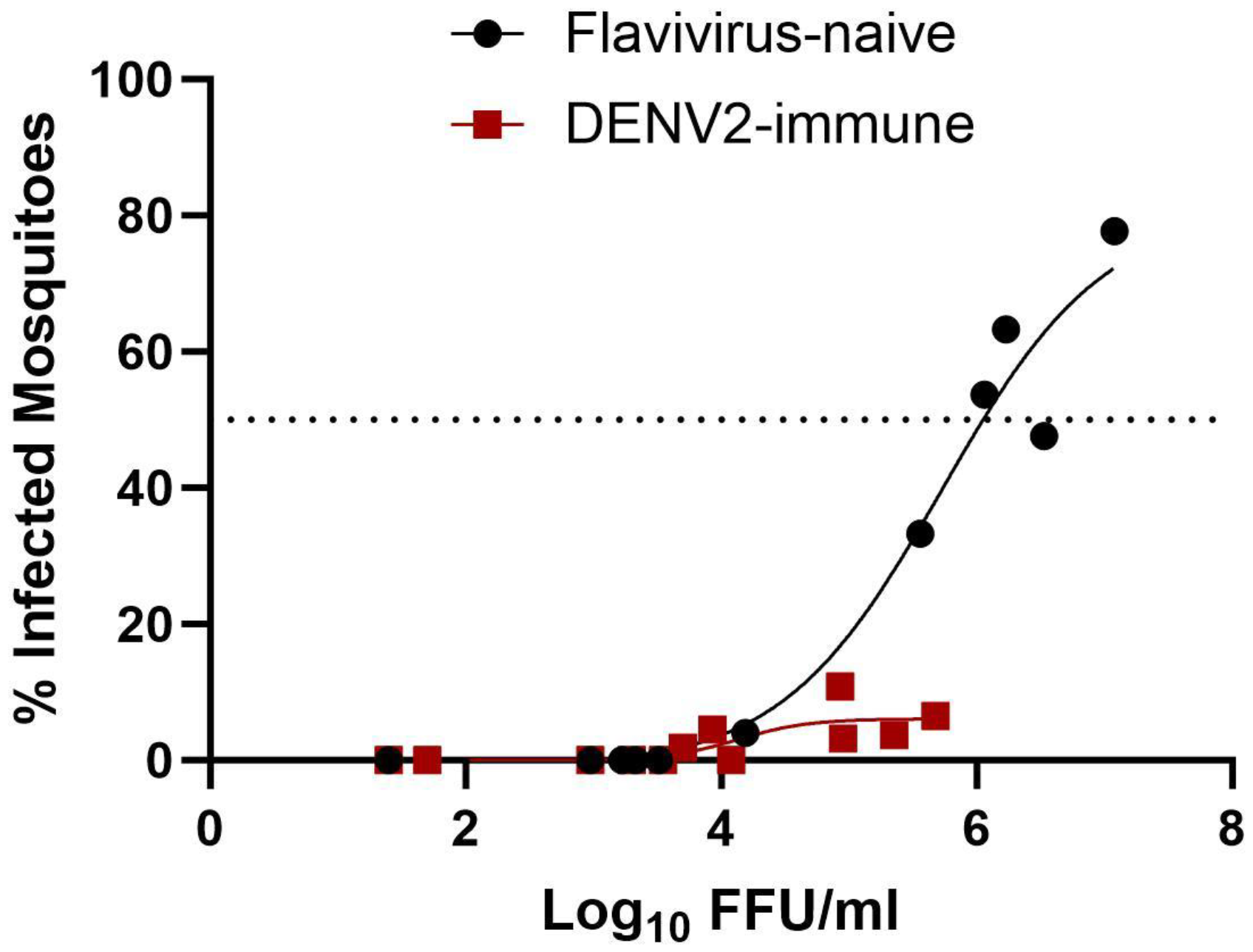
*Ae. aegypti* fed on DENV-immune, and YFV-challenged NHPs have low infection rates. Mosquito infectivity (%) was analyzed against NHP viremia for flavivirus-naïve and DENV-immune groups and curve-fitting using non-linear regression. Despite higher viremia at certain timepoints, most mosquitoes that fed on DENV-immune animals were resistant to infection.

### Heterologous immunity has mild effect on yellow fever severity

Upon inoculation with YFV, animals were observed daily for signs of illness. Temperatures were recorded via implanted loggers at 15-minute intervals, and weights and rectal temperatures were recorded from 1-5 and on 14 DPI and compared with baseline levels. No significant changes in weight (Figure 4a,b) or temperatures (Figure 4c,d) were detected among experimental groups. Blood tested for complete blood count (CBC) and serum chemistries revealed that most animals exhibited generalized leukopenia (22/23 animals) with specific populations of cells such as lymphocytes (23/23), neutrophils (22/23), and eosinophils (23/23) reduced from 2 to 5 DPI, compared to the baseline levels; leukopenia recovered by day 14 (Supplementary Figure 1,2). As a correlate of liver damage, alanine transaminase (ALT) and aspartate aminotransferase (AST) were measured, and high elevations were observed in flavivirus-naïve animals through day 14. In contrast, most DENV-2- (10/11) (Figure 4e, g) and ZIKV-immune (6/6) (Figure 4f, h) animals had elevated liver enzymes that resolved by day 14, when the levels for DENV-2-seronegative animals also resolved. The flavivirus-immune animals showed no significant changes in levels of alkaline phosphatase (ALP), creatine (CRE), blood urea nitrogen (BUN), or total bilirubin (TBIL) compared to -naïve animals (Supplementary Figure 3,4) up to day 5. After YFV infection, ZIKV- FRNT-negative but ELISA-positive animal FR1565 had a highly elevated neutrophil counts, BUN, CRE, TBIL (Supplementary Figure 2d, 4) and liver enzymes (ALP, ALT, and AST) (Figure 4f, h) on the day of euthanasia. Consistent with human infections exhibiting high viral load, neutrophils, elevated liver enzymes, and bilirubin associated with fatal YF (23), animal FR1565 exhibited all signs of fulminant fatal YF.

**Fig 4:**
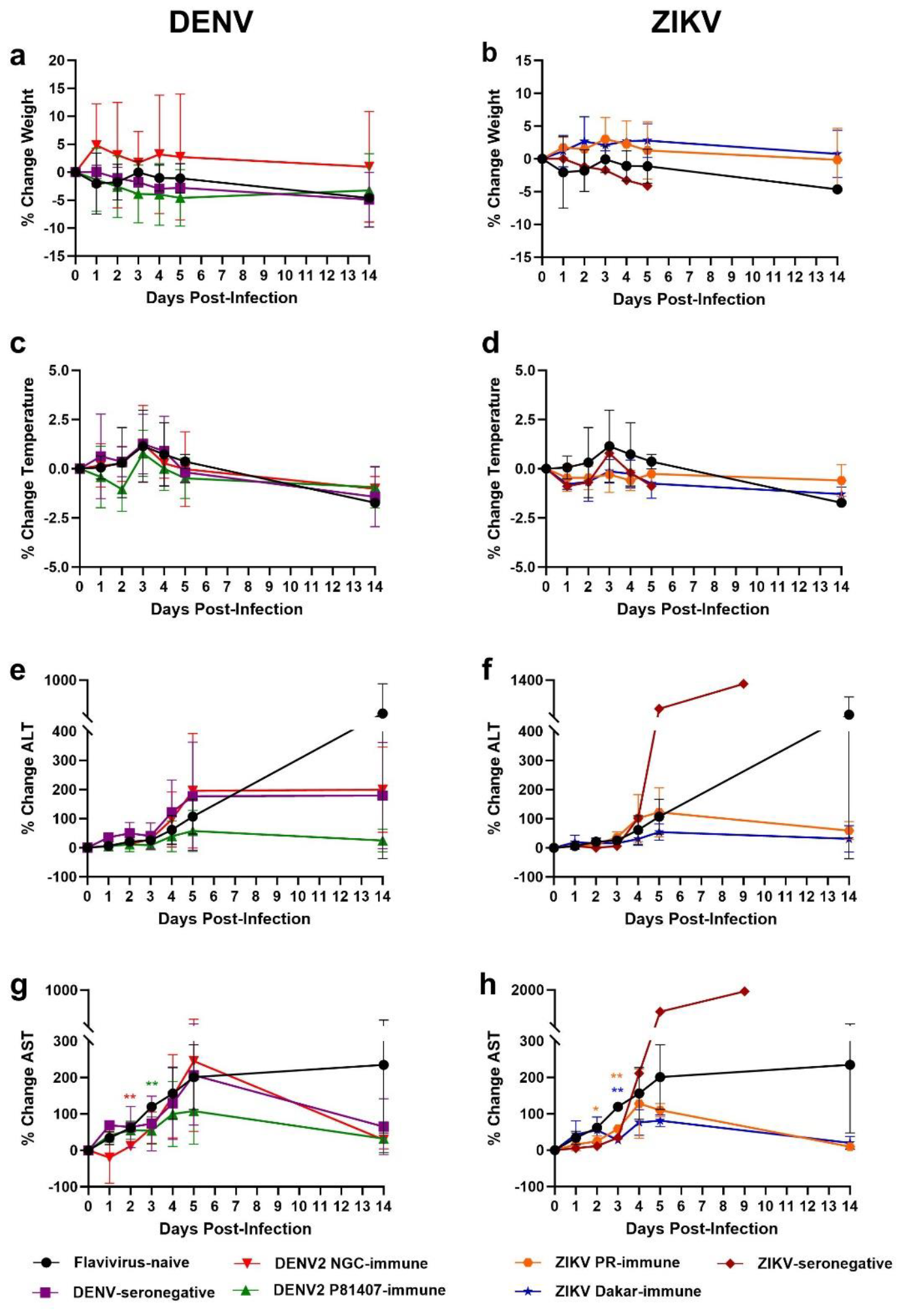
Prior flavivirus immunity does not affect the changes in clinical parameters. Animal weights **(4A, 4B)**, rectal temperatures **(4C, 4D)**, serum alanine transaminase (ALT) **(4E, 4F)**, and serum aspartate aminotransferase (AST) (**4G, 4H**) were measured from 1-5 and on 14 DPI and compared to baseline (day 0) values. Data were analyzed by two-way ANOVA with Dunnett’s multiple comparison test. The means of each group were compared with the flavivirus-naïve group at each time point. Statistical significance (*) color is attributed to the significant difference between that group compared to the flavivirus-naïve group. For example, in (4g), at 2DPI, flavivirus-naïve animals had a significantly higher change in AST compared to DENV-2 NGC-immune animals as shown by red^*^

### Heterologous immunity does not affect cytokine responses to YFV infection

The levels of certain cytokines and chemokines like interleukin-6 (IL-6), monocyte chemoattractant protein-1 (MCP-1), interferon-inducible protein (IP-10), tumor necrosis factor-α (TNF-α), and IL-1 receptor antagonist (IL-1RA) are significantly higher in fatal than non-fatal human YF *(24*). To understand the impact of heterologous flavivirus immunity on cytokine and chemokine responses, we conducted a Milliplex assay on 5 DPI samples and immediately prior to euthanasia [day 14, except day 9 for FR1565 (earlier euthanasia based on clinical scoring)] and compared them to baseline levels. Of 23 cytokines and chemokines measured, 18 remained below the limit of detection for all animals, except FR1565 (Figure 5a). IL1-ra is an anti-inflammatory cytokine that binds to the IL-1 receptor and suppresses pro-inflammatory cytokines like IL-1, TNF-α, and type I interferon. All 23 animals had elevated IL-1ra on day 5. At euthanasia, most animals had either reduced or maintained IL-1ra levels with the exception of FR1565, which had a 1674-fold increase compared to baseline (Figure 5b). Increased IL-1ra in the early stages of infection indicate a modulation of the inflammatory response. However, extremely high IL-1ra (as seen in FR1565) could affect the over-induction of anti-viral cytokines, contributing to the switch from a balanced immune environment to inflammation- induced tissue damage (*2*5). MCP-1 is a chemoattractant for monocytes as well as other immune cells, and can be secreted in response to activated pathogen recognition receptors, or by other cytokines like IL-6, IFN-β, and TNFα (*2*6). Levels of MCP-1 were elevated on day 5 in all animals, but declined by day 14 except in FR1565, which had a 4.8-fold increase compared to baseline levels (consistent with fatal human YF) (Figure 5c). Soluble CD40L (sCD40L) is primarily released by platelets upon viral activation, and high levels are associated with disease progression in HIV, DENV, and Influenza A (*27*). In hemorrhagic dengue, sCD40L levels are significantly decreased in patients with plasma leakage compared to those without, an indication of thrombocytopenia *(28*). Most of the 23 animals we infected with YFV had slight changes in sCD40L, except FR1565, which had a 42-fold decrease, suggesting destruction of platelets (Figure 5d). IL-8 (CXCL8) is a chemokine that plays a key role in inflammation by attracting neutrophils and T cells. For the majority of our animals, IL-8 remained unchanged or declined slightly after YFV infection, consistent with previous studies on macaques (*29*) (Figure 5e). Interestingly, IL-8 was elevated in ZIKV-immune animals. High levels of IL-8 have been reported in convalescent ZIKV patients (*30*), suggesting that prior ZIKV immunity could be an important factor in elevating IL-8 levels post-YFV-infection.

**Fig 5:**
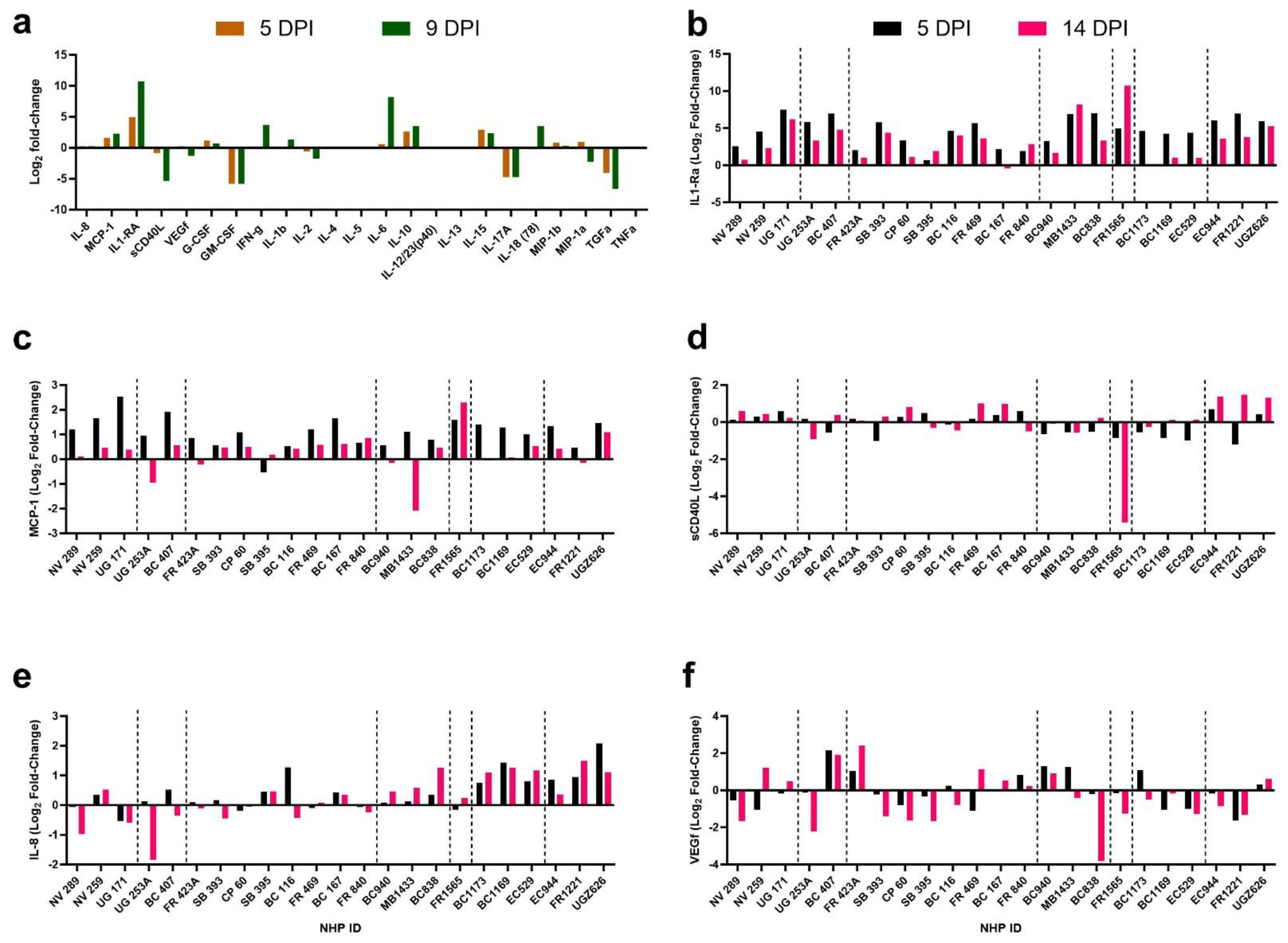
Prior heterologous immunity does not affect the cytokine response to YFV infection. Log_2_ fold-change in 23 different cytokines and chemokines measured in sera of FR1565 (ZIKV-exposed, seronegative animal) on days 5 and 9 post-YFV infection and compared with baseline levels **(5A)**. Log_2_ fold-change in IL1-ra **(5B)**, MCP-1 (**5C**), sCD40L (**5D**), IL-8 (**5E**), and VEGf (**5F**) at 5 DPI and euthanasia timepoints (14 DPI for all animals, 9 DPI for FR1565). Data shown for 23 animals; groups from left to right divided by dotted line are as followed: Flavivirus-naïve, DENV-2-seronegative, DENV-2 P81407-immune, DENV-2 NGC-immune, ZIKV-seronegative, ZIKV PRVABC59-immune, ZIKV DakAr-immune.

## Discussion

Mosquito-borne viruses infect hundreds-of-millions of people each year, with major health, economic, and societal impacts (*31*). The capacity of arboviruses to mutate and expand their host and geographic ranges, the projected impacts of climate change, the lack of vaccines and therapeutics, and the existence of immunologically susceptible populations, raise profound concerns regarding their continued emergence (*12*). Given that more than 2 billion individuals reside in Asian countries infested by *Ae. aegypti* (*31*), the potential consequences of YFV establishing itself in this region are grave, particularly since other mosquito-borne viruses like CHIKV and ZIKV arising from African enzootic cycles have repeatedly and efficiently established urban transmission cycles there *(12)*. Therefore, the reasons that YFV has not emerged into Asia in the same manner remain poorly understood.

An intriguing observation arose during the major 2016-2019 YFV outbreak in South America. Despite transmission in regions with large *Ae. aegypti* populations and limited vaccine coverage, no evidence was reported of the urban epidemic cycle involving this vector and human amplification. To explain this conundrum and the historic lack of YF in Asia, we investigated the potential role of immune barriers. We demonstrated that pre-existing immunity to DENV-2 or ZIKV in NHPs reduces YFV viremia, dramatically reducing mosquito infectivity. To ensure that our study reflected typical convalescent immunity and not an acute response, we allowed immunity to develop for at least 6-9 months post-primary infection. Overall, the hyperendemicity of DENV in Asia and South America, but lower seroprevalence in Africa, where urban *Ae. aegypti*-borne YFV outbreaks continue *(6,32*), further supports the hypothesis that flavivirus immunity, especially DENV, plays a major role in preventing urban YFV emergence in South America and Asia, but less so in Africa where seroprevalence is lower, by limiting the number of competent human amplifying hosts.

Intriguingly, we observed that infection of mosquitoes that fed on a DENV-2-immune animal was 20-30% lower compared to flavivirus-naïve animals with similar YFV viremia levels. This suggests that, in addition to directly reducing viremia, DENV-2 immunity may have a secondary blocking effect against YFV infection in mosquitoes. Mosquito-borne flaviviruses can utilize NS1 proteins produced in the serum to enhance oral infection mosquitoes *(33)*. Thus, it is possible that YFV NS1 is partially neutralized by heterologous NS1 antibodies, further reducing mosquito infection; this hypothesis deserves further attention.

Despite significant differences in mosquito infectivity, we did not observe a striking difference in YF disease outcomes in macaques, as indicated by serum chemistries, and cytokine levels. Since South American strains of YFV are milder compared to African strains in laboratory NHP models, we did not expect to see fatal outcomes in flavivirus-naïve or -immune animals *(34*). The elevated liver enzyme levels in most -immune animals resolved by day 14, suggesting faster recovery and less liver damage. In ZIKV-FRNT-negative but ELISA-positive animal FR1565, all clinical findings recapitulated fatal human YF. The low levels of ZIKV non-neutralizing IgG in this animal suggested the possibility of antibody-dependent enhancement, as has been reported in sequential flavivirus infections (mainly DENV) (*35)*. However, with only one animal showing this outcome, we cannot conclude that low-level ZIKV immunity exacerbated the YF outcome, and further studies are needed.

While our goal was to investigate the role of heterologous flavivirus immunity on urban YF emergence, additional work is needed to determine the impacts of various levels of pre-existing flavivirus immunity on YF severity, including additional intervals between sequential infections. Of the 720 mosquitoes from which we harvested saliva at 3 and 4 DPI, only one had detectable YFV. Due to restrictions on the number of mosquitoes that can be exposed to each animal, and to maximize the sample size, we harvested mosquitoes at a single timepoint. For future work, multiple timepoints, including greater periods of extrinsic incubation, should further address the transmission potential. However, relatively low YFV transmission efficiency of *Ae. aegypti* populations, even those clearly incriminated in epidemics, is not uncommon (*36*).

We did not determine the mechanism of cross-protection in flavivirus-immune NHPs, other than to rule out cross-neutralizing antibodies. The role of both humoral and cell-mediated immunity should be investigated in murine models, as well as the potential role of flavivirus NS1 antibodies in blocking mosquito infections to limit transmission. Also, it is important to investigate the implications for reciprocal effects on DENV and ZIKV infection following YF vaccination or infection. Although our central hypothesis reflected the assumption that human flavivirus immunity, especially for DENV and ZIKV, is lower in Africa than in Asia or South America (*32,37–40*), more data on seroprevalence in locations such as Angola, which recently experienced urban YF, would be useful. In addition, further studies utilizing African strains of YFV and African *Ae. aegypti* mosquitoes could expand the implications of our work. Finally, although primarily a rural disease (*41,42)*, possible cross-protective immunity from Japanese encephalitis (JE), which is endemic in Asia, or from JE vaccination, should also be studied.

In summary, we demonstrated that prior immunity in DENV- or ZIKV-endemic regions likely plays a major role in preventing urban YFV emergence by reducing the population of amplification-competent people who can infect *Ae. aegypti* mosquitoes. These findings have major implications for future YF risk as flaviviruses continue to spread due to anthropogenic changes (*43*). Moreover, the expansion of *Ae. aegypti* due to climate change will increase the risk of YF in areas such as Southern USA with limited flavivirus immunity that have not experienced epidemics in over a century (*43*).

## Materials and Methods

No statistical methods were used to predetermine the sample size as the NHPs were originally utilized in a separate experiment in keeping with the ethical goal of reducing the total number of animals utilized in research. The experiments were not randomized. The investigators were not blinded to allocation during experiments and outcome assessment.

### Animals, Study Design, and Ethics Statement

Twenty-three healthy adolescent cynomolgus macaques (11 males and 12 females, 2.5-5.5 years old, 3-5.5 kg) were transferred from a previous study, where they were exposed to mosquitoes infected with PBS (3/23), DENV-2 NGC (3/23), DENV-2 P8-1407 (10/23), ZIKV PRVABC59 (4/23), and ZIKV DakAr 41525 (3/23). After primary exposure, animals transferred to our study were allowed to build immunity for 6-9 months. All animals were then inoculated subcutaneously with 0.5ml of YFV BR17 at a final dose of 1 x 104 FFU.

Temperatures were recorded at 15-minute intervals on loggers (Star Oddi, Iceland) placed in the abdominal cavity. Animals were fed twice a day with commercial chow. For enrichment, animals were fed an additional item like fruit or vegetable. Health checks were performed at least twice daily from the day of infection until the end of the study. Body weight and rectal temperatures were measured on study days 1-5 and 14. On specified days, animals were anesthetized by intramuscular injection with Ketamine (5-20mg/kg). Blood samples were collected into serum separator tubes (SST, Greiner Bio-one) and tubes containing ethylenediaminetetraacetic acid (K2-EDTA, BD Vacutainer). Whole blood was analyzed for various parameters using a HEMAVET 950 multispecies hematology instrument (Drew Scientific, Miami, FL). Serum was analyzed using a Preventative Care Profile Plus rotor on a VetScan VS2 Chemistry Analyzer (Abaxis, Union City, CA). A cardboard carton with a mesh top containing 100 female *Ae. aegypti* (mosquitoes from Salvador, Brazil, maintained for 18 generations at the UTMB Insectary) were allowed to feed on NHP ears for 5-15 minutes.

Animals were euthanized at the end of the study by pentobarbital overdose. All experiments were performed in full compliance with the guidelines established by the Animal Welfare Act for the housing and care of laboratory animals and conducted as laid out in the University of Texas Medical Branch Institutional Animal Care and Use Committee (UTMB-IACUC) approved protocol (protocol #2101002, approved 21 January 2021; protocol #2208052, approved 02 September 2022). UTMB is an Association for Assessment and Accreditation of Laboratory Animal Care International-accredited facility.

### Cells

African green monkey kidney epithelial (Vero E6) and Ae. albopictus (C6/36) cells were provided by the World Reference Center for Emerging Viruses and Arboviruses (WRCEVA). Vero E6 cells were grown in minimal essential medium (MEM, Gibco, Grand Island, NY) supplemented with 10% fetal bovine serum (FBS, Atlanta Biologics, Atlanta, GA), 1% penicillin streptomycin (PennStrep,Gibco, Grand Island, NY), 1% sodium bicarbonate (Gibco, Grand Island, NY), and 1% glutamax (Gibco, Grand Island, NY). C6/36 cells were grown in MEM and L-15 media at a ratio of 1:1, supplemented with 10% FBS, 1% PennStrep, 1% glutamax, 1% sodium bicarbonate, 10% tryptose phosphate broth solution (TPB, Sigma, St Louis, MO), 2% non-essential amino acid solution (Sigma, St Louis, MO). Cells were maintained at 37°C with 5% CO2.

### Viruses

The YFV BR17 stock was generated at the WRCEVA (originally acquired from Dr. Betânia Drumond, Universidade Federal de Minas Gerais, Brazil). The virus was isolated from an infected Alouatta guariba in 2017 in Minas Gerais during the YF epidemic. The virus was passaged three times in C6/36 cells (Ae. albopictus cell line) to generate a low passage contemporary YFV BR17 stock that was utilized for the primate challenge.

Stocks of DENV1 16007, DENV3 D83-144, DENV4 703-4, DENV-2 NGC, DENV-2 P8- 1407, ZIKV PRVABC59, and ZIKV DakAr 41525 were acquired from WRCEVA and passaged once in C6/36 cells (for DENV-2 strains) and Vero E6 (for ZIKV strains). These viral stocks were used for foci reduction neutralization test (FRNT).

### Mosquito Feeding Assay

*Aedes aegypti* obtained from Salvador, Brazil, were maintained in the UTMB Insectary and utilized at generation F18. Female mosquitoes (100 in each cup) were sucrose-starved for approximately 16 h. On days 1-5 post-YFV infection, one separate cup of mosquitoes was exposed to individual NHPs on the ear(s) for 5-15 minutes, to make a total of five separate cups per animal (one per day) and 115 cups for the entire study of 23 animals. Fully engorged mosquitoes were maintained at 28°C with a 16-hour light/8-hour dark cycle with ad libitum access to 10% sucrose. At 14 days post-feeding, legs and wings were removed from cold-anesthetized mosquitoes and collected in 0.5ml homogenization media (DMEM supplemented with 2% FBS, 1% antibiotic-antimycotic) with 5mm stainless steel sterile bead (Glen Mills, Clifton, NJ). Mosquitoes were force-salivated by inserting the proboscis in a 10μl pipette tip containing 10μl FBS for approximately 30 minutes. Saliva-containing FBS was then collected in 0.1ml homogenization media and the remaining mosquito body was collected in 0.5ml homogenization media. All samples were stored at -80°C until the day of processing. Prior to the focus-forming assay to determine the titer and infectivity, samples were thawed and homogenized at a frequency of 26/s for 1 minute in a TissueLyser II (Qiagen, Germantown, MD) and centrifuged at 16,500xg for 5 minutes. Unfortunately, due to technical difficulties with an incubator that prevented the mosquitoes from surviving for the full 14 days post-feed, we could not complete the harvests of mosquitoes that fed on 5 DENV-2 P81407 immune NHPs (NHP ID: UG253A, FR 423A, SB 393, CP 60, SB 395). Therefore, mosquito assay data is only available for 18 animals.

### Focus-Forming Assays

Vero E6 cells were seeded in each well of 12 well or 96 well plates and cultured at 37°C, 5% CO2 for 24 h until 95% confluency. To determine the viremia, serum samples were serially diluted with Vero maintenance media (MEM, 2% FBS, 1% Penn-Strep, 1% Glutamax, 1% sodium bicarbonate) and transferred to the pre-seeded Vero E6 cells in 12 well plates. The samples were incubated at 37°C for 1 h. After incubation, 1ml of overlay media (Opti-MEM (Gibco, Grand Island, NY) supplemented with 1% carboxymethyl cellulose (Sigma Aldrich, St. Louis, MO), 2% FBS, 1% Penn-Strep was added to each well. After a 5-day incubation, formalin was added to fix the infected plates. Plates were washed thrice with 1X DPBS (Gibco, Grand Island, NY), followed by blocking with 1ml of non-fat dry milk (Apex Chemicals and Reagents, Houston, TX) in 1X DPBS (5% w/v) for 30 minutes on a plate rocker. After discarding the blocking buffer, anti-YFV primary antibody (mouse immune ascitic fluid acquired from WRCEVA) diluted at 1:1,000 in block buffer was used for staining overnight at a volume of 300μl per well. The following day, plates were washed (3x with 1X DPBS) followed by adding the secondary antibody (affinity purified antibody peroxidase labeled goat anti-mouse IgG, SeraCare, Milford, MA) diluted at 1:2,000 in block buffer at a volume of 300μl per well and incubated on the plate rocker for 1 h. Following a final three washes, plates were stained using KPL TrueBlue Peroxidase Substrate (SeraCare, Milford, MA) at a volume of 150μl per well, and foci were counted to determine the final infectious viral load expressed as FFU/mL. For mosquito titrations, homogenized samples were titrated on 96-well plates with Vero E6 cells, fixed at day 4, and stained with 40μl primary and secondary antibodies and 25μl TrueBlue. Foci were manually counted from images obtained on Cytation 7 plate reader (Agilent Life Sciences, Santa Clara, CA). For mosquito infectivity, 100μl of homogenate was used to infect Vero E6 cells in 96-well plates without dilution, fixed at day 7, and stained as with mosquito titrations.

### Foci Reduction Neutralization Test (FRNT)

To determine the neutralizing capacity of antibodies in sera, FRNTs were conducted on baseline samples (prior to YFV infection). Samples from primates that were exposed to DENV-2 NGC or P8-1407 were tested against all four serotypes of DENV (using the strains mentioned above) and YFV (BR17). Samples from animals exposed to ZIKV PRVABC59 or DakAr 41525 were tested against ZIKV and YFV. Briefly, sera were serially diluted two-fold and mixed with 800 FFU of the respective virus. Serum-virus mixture was incubated at 37°C for 1 h followed by infection of Vero E6 cells on 12 well plates. Samples tested against DENV1, 3, and 4 were fixed on day 4, and DENV-2 and YFV were fixed on day 5, and ZIKV was fixed on day 3. Plates were stained as mentioned above with the respective primary antibodies. The neutralizing titer was represented as the reciprocal of the highest dilution of serum that inhibited 50% of foci (FRNT50). Samples that scored below the limit of detection (<1:20) were considered seronegative.

### Cytokine and Chemokines

Sera samples collected on day 5 and on the euthanasia day (14 or 9), were analyzed for 23 cytokines and chemokines using the Milliplex NHP Cytokine Magnetic Bead Panel (Millipore Sigma, Burlington, MA) following the manufacturer’s protocol. Results were analyzed on Bio-Plex 200 (Biorad, Hercules, CA). Standard curves of known concentrations of cytokines were used to convert fluorescence units into cytokine concentration units (pg/mL).

### Statistical Analysis

Descriptive statistics have been provided in figure legends. Log10 transformed viremia titers in each group were analyzed by two-factor repeated measures ANOVA with Tukey’s multiple comparison test. Mosquito infection rates were analyzed by two-tailed Fisher’s exact test by combining the data in each group and comparing with the flavivirus-naïve group. Log10 transformed viral titers in mosquito groups were compared with flavivirus-naïve group by one- way ANOVA with Dunnet’s multiple comparison test. Samples that had viremia and viral loads below the limit of detection were graphed as ½ of LOD and the same value was used for statistical analyses. For comparisons between more than two groups and more than two timepoints, analyses by two-way ANOVA repeated measures and appropriate post-hoc tests were conducted. All statistical analysis and graphing was performed with GraphPad Prism Version 10.

## Acknowledgments

We thank the Animal Resource Center at UTMB for their assistance with primate work, and the Insectary staff (Ruimei Yun, Jiehua Zhou) at UTMB for providing mosquitoes from the colony.

## Funding

World Reference Center for Emerging Viruses and Arboviruses, National Institute of Health grant R24AI120942 (SCW)

Centers for Research in Emerging Infectious Diseases (CREID) (SCW)

West African Center for Emerging Infectious Diseases (WAC-EID) U01-AI151801 (SCW)

The Coordinating Research on Emerging Arboviral Threats Encompassing the Neotropics (CREATE-NEO) U01 AI151807 (NV)

National Institute of Health R01 AI145918 (NV)

National Institute of Health K99AI168484 (RKC)

National Council for Scientific and Technological Development (CNPq)/Brazil (BPD)

Center for Vector-borne and Zoonotic Diseases, University of Texas Medical Branch (DPS)

Jeane B. Kempner Fellowship, University of Texas Medical Branch (DPS)

## Author contributions

Conceptualization: DPS, KSP, JAP, SCW

Methodology: DPS, KSP, JAP, SCW

Investigation: DPS, JAP, DS, BM, JW, SRA, RKC, LS, BD, NV, KSP, SCW

Visualization: DPS, KSP, JAP, SCW

Funding acquisition: NV, SCW

Writing-original draft: DPS, SCW

Writing-review and editing: DPS, JAP, DS, BM, JW, SRA, RKC, LS, BD, NV, KSP, SCW

## Data and materials availability

All data are available in the main text or the supplementary materials.

