## Supplementary Figures and Tables for "Yellow Fever Emergence: Role of Heterologous Flavivirus Immunity in Preventing Urban Transmission"

**The PDF file includes:**

Figs. S1 to S4
Tables S1-S3, and S5
Table S4 is available separately as .xlsx file due to the size

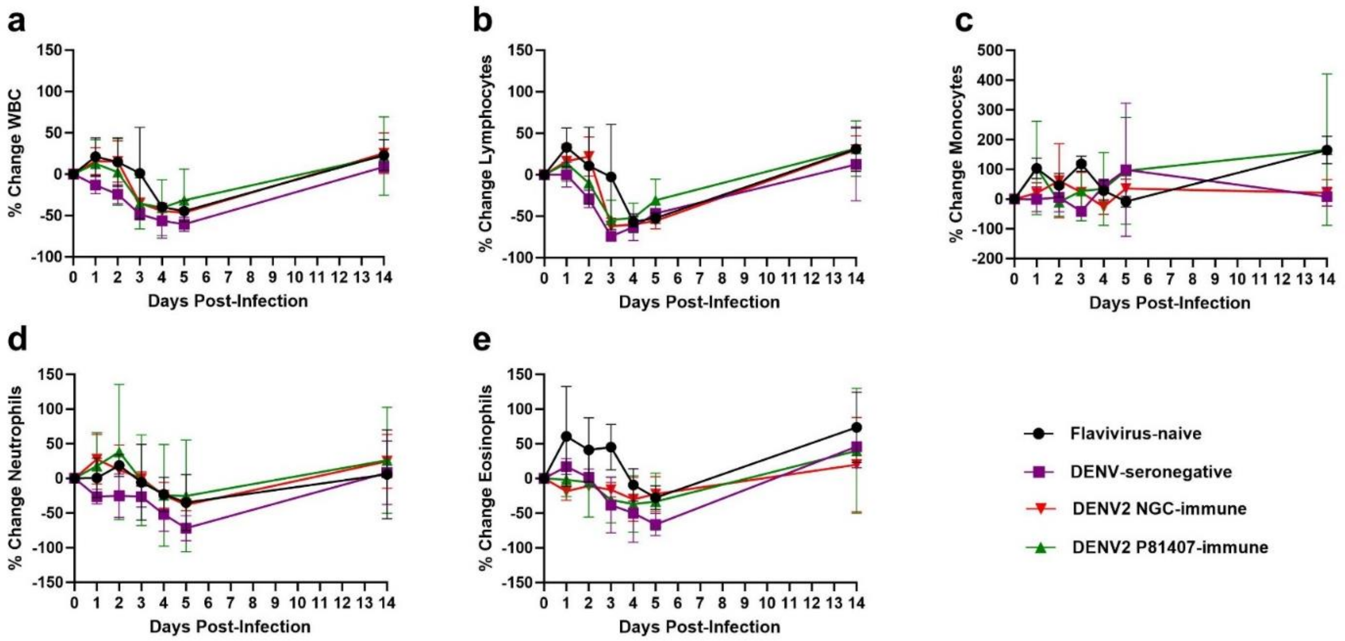


**Fig. S1. Percentage change in WBCs, lymphocytes, monocytes, neutrophils, and eosinophils** Values measured on 1-5 and on 14 DPI compared with baseline levels in flavivirus-naïve and DENV-2-exposed NHPs. Each group was compared with the other by two-way ANOVA repeated measures. No significant difference was observed in any of the groups.

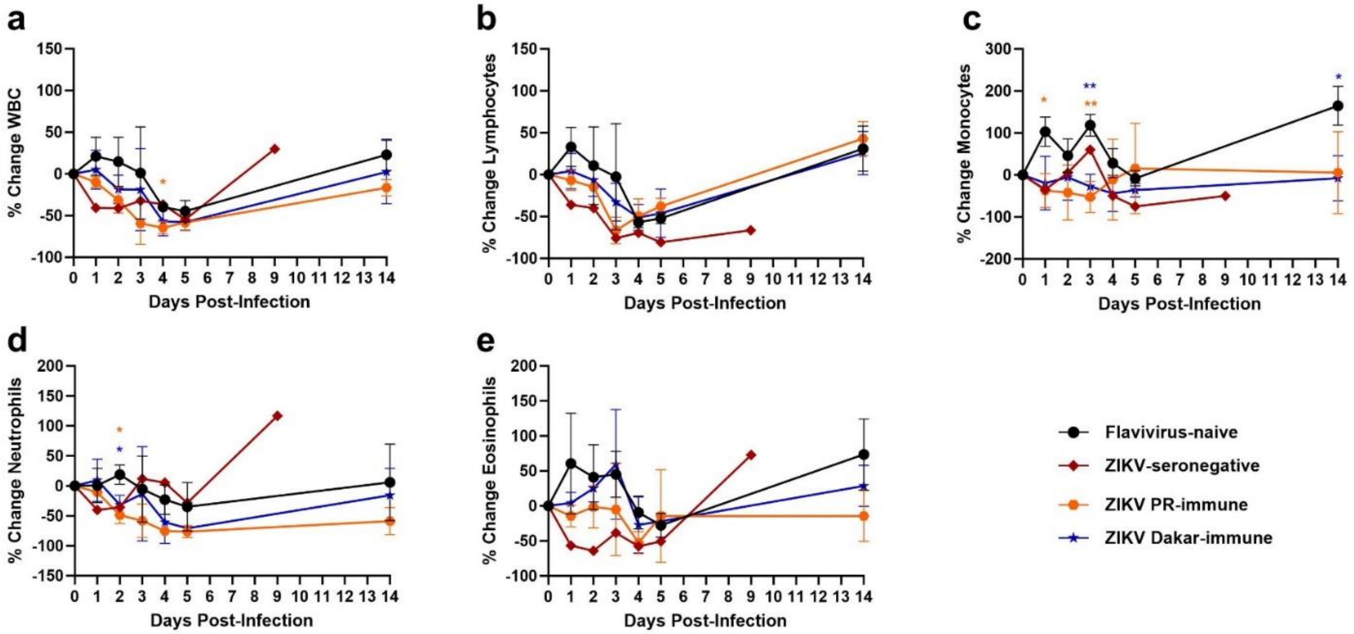


**Fig. S2. Percentage change in WBCs, lymphocytes, monocytes, neutrophils, and eosinophils** Values measured on 1-5 and 14 DPI compared with baseline levels in flavivirus-naïve and ZIKV-exposed NHPs. Each group was compared with the other by two-way ANOVA repeated measures. Since the ZIKV-seronegative group has only one animal, this group was excluded from analyses. Statistical significance (*) color is attributed to the significant difference between that group compared to the flavivirus-naïve group. *p<0.05, ** p<0.01.

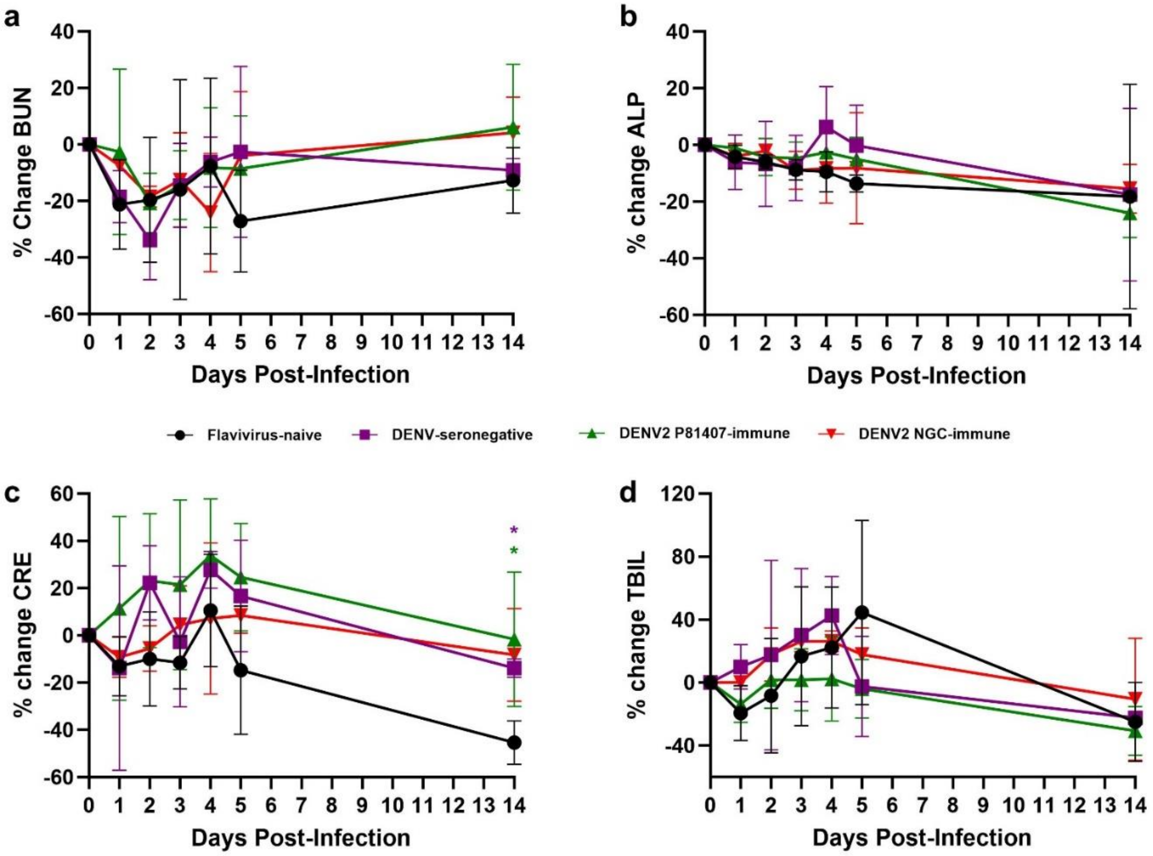


**Fig. S3. Percentage change in blood urea nitrogen (BUN), alanine phosphatase (ALP), creatine (CRE), total bilirubin (TBIL)**

Values measured on 1-5 and 14 DPI compared with baseline levels in flavivirus-naïve and DENV-2-exposed NHPs. Each group was compared with the flavivirus-naïve group by two-way ANOVA repeated measures. Statistical significance (*) color is attributed to the significant difference between that group compared to the flavivirus-naïve group. *$p<0.05$.

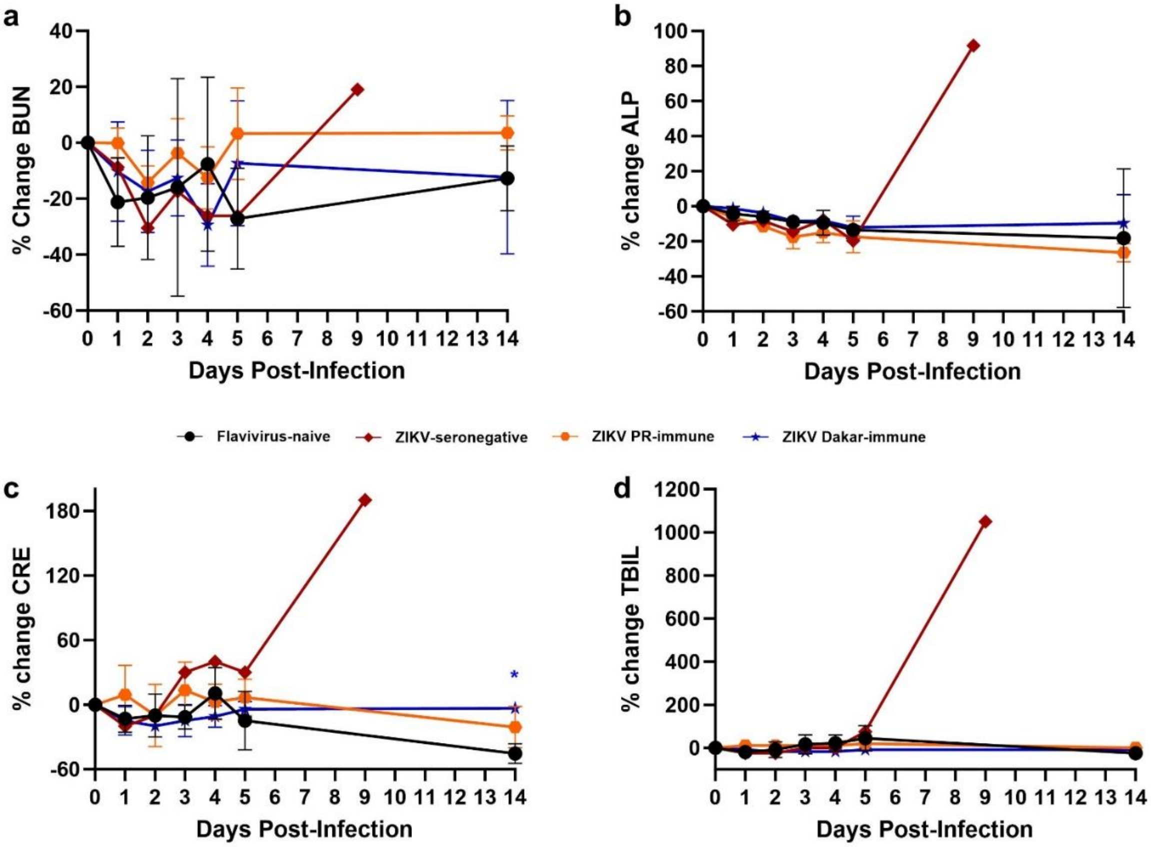


**Fig S4. Percentage change in blood urea nitrogen (BUN), alanine phosphatase (ALP), creatine (CRE), total bilirubin (TBIL)**

Values measured on 1-5 and 14 DPI compared with baseline levels in flavivirus-naïve and ZIKV-exposed NHPs. Each group was compared with the flavivirus-naïve group by two-way ANOVA repeated measures. Since the ZIKV-seronegative group has only one animal, this group was excluded from analyses. Statistical significance (*) color is attributed to the significant difference between that group compared to the flavivirus-naïve group. *$p<0.05$.

**Table S1. FRNT$_{50}$ measured on Day 0 before YFV challenge.**

| NHP ID | Primary exposure | FRNT$_{50}$ (1/n) | | | | | |
|---|---|---|---|---|---|---|---|
| | | YFV | DENV1 | DENV-2 | DENV3 | DENV4 | ZIKV |
| NV 289 | Naïve | <20 | <20 | <20 | <20 | <20 | <20 |
| NV 259 | Naïve | <20 | <20 | <20 | <20 | <20 | <20 |
| UG 171 | Naïve | <20 | <20 | <20 | <20 | <20 | <20 |
| UG 253A | DENV-2 P81407 | <20 | <20 | <20 | <20 | <20 | # |
| BC 407 | DENV-2 P81407 | <20 | <20 | <20 | <20 | <20 | # |
| FR 423A | DENV-2 P81407 | <20 | <20 | 2560 | <20 | <20 | # |
| SB 393 | DENV-2 P81407 | <20 | <20 | 320 | <20 | <20 | # |
| CP 60 | DENV-2 P81407 | <20 | <20 | 40 | <20 | <20 | # |
| SB 395 | DENV-2 P81407 | <20 | <20 | 640 | <20 | <20 | # |
| BC 116 | DENV-2 P81407 | <20 | <20 | 80 | <20 | <20 | # |
| FR 469 | DENV-2 P81407 | <20 | <20 | 640 | <20 | <20 | # |
| BC 167 | DENV-2 P81407 | <20 | <20 | 1280 | <20 | <20 | # |
| FR 840 | DENV-2 P81407 | <20 | <20 | 640 | <20 | <20 | # |
| BC940 | DENV-2 NGC | <20 | <20 | 640 | <20 | <20 | # |
| MB1433 | DENV-2 NGC | <20 | <20 | 320 | <20 | <20 | # |
| BC838 | DENV-2 NGC | <20 | <20 | 640 | <20 | <20 | # |
| FR 1565 | ZIKV PRVABC59 | <20 | # | | | | <20 |
| BC 1173 | ZIKV PRVABC59 | <20 | # | | | | 320 |
| BC 1169 | ZIKV PRVABC59 | <20 | # | | | | 160 |
| EC 529 | ZIKV PRVABC59 | <20 | # | | | | 2560 |
| EC 944 | ZIKV DakAr 41525 | <20 | # | | | | 640 |
| FR 1221 | ZIKV DakAr 41525 | <20 | # | | | | 640 |
| UG 2626 | ZIKV DakAr 41525 | <20 | # | | | | 640 |

### DENV-2 and ZIKV-exposed samples were not evaluated against each other as it is beyond the scope of this manuscript

**Table S2. Log$_{10}$-transformed YFV viremia on days 1-5 post-infection.**

| Primary exposure | FRNT$_{50}$ against DENV-2 (1/n) | NHP ID | YFV Viremia (Log$_{10}$ FFU/mL) Days Post-Infection | | | | |
|---|---|---|---|---|---|---|---|
| | | | 1 | 2 | 3 | 4 | 5 |
| **PBS** | <LOD | NV 289 | <1.70 | 3.22 | 5.56 | 6.23 | 3.51 |
| | <LOD | NV 259 | <1.70 | <1.70 | 2.98 | 4.19 | <1.70 |
| | <LOD | UG 171 | <1.70 | 3.33 | 6.06 | 7.08 | 6.53 |
| **DENV-2 P81407** | <LOD | UG 253A | <1.70 | 2.92 | 6.02 | 5.04 | <1.70 |
| | <LOD | BC 407 | <1.70 | 3.15 | 5.39 | 5.32 | <1.70 |
| | 2560 | FR 423A | <1.70 | <1.70 | <1.70 | <1.70 | <1.70 |
| | 320 | SB 393 | <1.70 | <1.70 | 4.89 | 4.68 | <1.70 |
| | 40 | CP 60 | <1.70 | 3.32 | 5.02 | 6.28 | <1.70 |
| | 640 | SB 395 | <1.70 | <1.70 | <1.70 | <1.70 | <1.70 |
| | 80 | BC 116 | <1.70 | <1.70 | 4.95 | <1.70 | <1.70 |
| | 640 | FR 469 | <1.70 | <1.70 | 3.30 | 4.08 | <1.70 |
| | 1280 | BC 167 | <1.70 | <1.70 | <1.70 | <1.70 | <1.70 |
| | 640 | FR 840 | <1.70 | <1.70 | <1.70 | <1.70 | <1.70 |
| **DENV-2 NGC** | 640 | BC940 | † | 1.70 | 3.54 | 3.93 | <1.70 |
| | 320 | MB1433 | † | 1.70 | 3.70 | 4.93 | <1.70 |
| | 640 | BC838 | † | 2.98 | 5.35 | 5.68 | <1.70 |

Limit of detection (LOD) is 50 FFU/mL (1.7 log$_{10}$ FFU/mL)

**Table S3. Mosquito infection rate at peak YFV viremia timepoint**

| Primary exposure | $FRNT_{50}$ against ZIKV (1/n) | NHP ID | YFV Viremia ($Log_{10}$ FFU/mL) Days Post-Infection | | | | |
|---|---|---|---|---|---|---|---|
| | | | 1 | 2 | 3 | 4 | 5 |
| **PBS** | <LOD | NV 289 | <1.70 | 3.22 | 5.56 | 6.23 | 3.51 |
| | <LOD | NV 259 | <1.70 | <1.70 | 2.98 | 4.19 | <1.70 |
| | <LOD | UG 171 | <1.70 | 3.33 | 6.06 | 7.08 | 6.53 |
| **ZIKV PRVABC59** | <LOD | FR1565 | † | 1.81 | 5.83 | 7.30 | 7.52 |
| | 320 | BC1173 | † | 3.51 | 4.98 | 4.29 | <1.70 |
| | 160 | BC1169 | † | <1.70 | 3.10 | 4.00 | <1.70 |
| | 2560 | EC529 | † | 1.70 | 3.65 | 2.68 | <1.70 |
| **ZIKV DakAr 41525** | 640 | EC944 | † | <1.70 | <1.70 | 2.10 | <1.70 |
| | 640 | FR1221 | † | 1.88 | 4.15 | 4.24 | <1.70 |
| | 640 | UGZ626 | † | <1.70 | 3.54 | 2.30 | <1.70 |

**† indicates the samples that were lost due to inconsistent storage temperatures**

**Table S5. Infection rate of mosquitoes analyzed by two-tailed Fisher's Exact Test**

| Timepoint | Group 1 | Group 2 | Significance? | p-value |
|---|---|---|---|---|
| 3 DPI | Flavivirus-naïve | DENV-2 NGC-immune | Yes | <0.0001 |
| | Flavivirus-naïve | DENV-2 P8-immune | Yes | <0.0001 |
| | Flavivirus-naïve | DENV-2-seronegative | No | 0.2678 |
| 4 DPI | Flavivirus-naïve | DENV-2 NGC-immune | Yes | <0.0001 |
| | Flavivirus-naïve | DENV-2 P8-immune | Yes | <0.0001 |
| | Flavivirus-naïve | DENV-2-seronegative | Yes | 0.0026 |
| 3 DPI | Flavivirus-naïve | ZIKV PR-immune | Yes | <0.0001 |
| | Flavivirus-naïve | ZIKV DakAr-immune | Yes | <0.0001 |
| | Flavivirus-naïve | ZIKV-seronegative | No | 0.0132 |
| 4 DPI | Flavivirus-naïve | ZIKV PR-immune | Yes | <0.0001 |
| | Flavivirus-naïve | ZIKV DakAr-immune | Yes | <0.0001 |
| | Flavivirus-naïve | ZIKV-seronegative | No | 0.3132 |
