## Supplementary Table 4 for "Yellow Fever Emergence: Role of Heterologous Flavivirus Immunity in Preventing Urban Transmission"

**Table S4. Mosquito infection rate at peak YFV viremia timepoint**

| Prior flavivirus immunity | NHP ID | Days Post-Infection | YFV Viremia (log$_{10}$ FFU/ml) | Total Infected Bodies (x) | Total Engorged (y) | Infection Rate = x/y *100) | Dissemination Rate = Total Infected Legs/x *100 |
|---|---|---|---|---|---|---|---|
| None | NV 289 | 3 | 5.56 | 10 | 30 | 33.33 | 0 (0) |
| | | 4 | 6.23 | 14 | 22 | 63.63 | 9 (64.3) |
| | NV 259 | 3 | 2.98 | 0 | 59 | 0 | † |
| | | 4 | 4.19 | 2 | 50 | 4 | 0 (0) |
| | UG 171 | 3 | 6.06 | 29 | 54 | 53.7 | 6 (20.7) |
| | | 4 | 7.08 | 49 | 63 | 77.77 | 4 (8.2) |
| DENV-2-seronegative | UG253A | 3 | 6.02 | | NA* | | |
| | | 4 | 5.04 | | NA | | |
| | BC 407 | 3 | 5.39 | 21 | 71 | 29.5 | 1 (4.8) |
| | | 4 | 5.32 | 18 | 69 | 24.8 | 2 (11.11) |
| DENV-2 P8 1407 | FR 423A | 3 | 1.4 | | NA | | |
| | | 4 | 1.4 | | NA | | |
| | SB 393 | 3 | 4.89 | | NA | | |
| | | 4 | 4.68 | | NA | | |
| | CP 60 | 3 | 5.02 | | NA | | |
| | | 4 | 6.28 | | NA | | |
| | SB 395 | 3 | 1.4 | | NA | | |
| | | 4 | 1.4 | | NA | | |
| | BC 116 | 3 | 4.95 | 1 | 31 | 3.2 | 0 (0) |
| | | 4 | 1.4 | 0 | 54 | 0 | † |
| | FR 469 | 3 | 3.3 | 0 | 75 | 0 | † |
| | | 4 | 4.08 | 0 | 73 | 0 | † |
| | BC 167 | 3 | 1.4 | 0 | 53 | 0 | † |
| | | 4 | 1.4 | 0 | 73 | 0 | † |
| | FR 840 | 3 | 1.4 | 0 | 50 | 0 | † |
| | | 4 | 1.4 | 0 | 59 | 0 | † |
| DENV-2 NGC | BC 940 | 3 | 3.54 | 0 | 73 | 0 | † |
| | | 4 | 3.93 | 3 | 66 | 4.5 | 1 (33.33) |
| | MB 1433 | 3 | 3.7 | 1 | 52 | 1.9 | 0 (0) |
| | | 4 | 4.93 | 6 | 55 | 10.9 | 0 (0) |
| | BC 838 | 3 | 5.35 | 1 | 27 | 3.7 | 0 (0) |
| | | 4 | 5.68 | 3 | 46 | 6.5 | 0 (0) |
| ZIKV PR-seronegative | FR 1565 | 3 | 5.83 | 7 | 44 | 9 | 2 (28.5) |
| | | 4 | 7.3 | 18 | 30 | 60 | 4 (22.2) |
| ZIKV PRVABC59 | BC 1173 | 3 | 4.98 | 1 | 27 | 3.7 | 0 (0) |
| | | 4 | 4.29 | 0 | 36 | 0 | † |
| | BC 1169 | 3 | 3.1 | 0 | 56 | 0 | † |
| | | 4 | 4 | 0 | 59 | 0 | † |
| | EC 529 | 3 | 3.65 | 0 | 67 | 0 | † |
| | | 4 | 2.68 | 0 | 41 | 0 | † |
| ZIKV DakAr 41525 | EC 944 | 3 | 1.4 | 0 | 59 | 0 | † |
| | | 4 | 2.1 | 0 | 69 | 0 | † |
| | FR 1221 | 3 | 4.15 | 0 | 27 | 0 | † |
| | | 4 | 4.24 | 0 | 17 | 0 | † |
| | UG 2626 | 3 | 3.54 | 0 | 57 | 0 | † |
| | | 4 | 2.3 | 0 | 49 | 0 | † |

*NA refers to groups of mosquitoes that did not survive for 14 days post-feeding due to technical issues with the incubator.

Dissemination rate (DR) refers to the number of positive mosquito legs/ total number of mosquitoes infected. Legs from mosquitoes that were positive for infection were utilized to measure dissemination. † refers to the samples that were not assayed due to negative infection in bodies
